# A hacked kitchen scale-based system for quantification of grip strength in rodents

**DOI:** 10.1101/2020.07.23.217737

**Authors:** Jan Homolak, Davor Virag, Ivan Kodvanj, Ivica Matak, Ana Babic Perhoc, Ana Knezovic, Jelena Osmanovic Barilar, Vladimir Trkulja, Melita Salkovic-Petrisic

## Abstract

Assessment of neuromuscular function is critical for understanding pathophysiological changes related to motor system dysfunction in many rodent disease models. Among methods used for quantification of grip performance in rodents, gauge-based grip strength meters provide the most reliable results, however, such instruments are unaffordable by many laboratories. Here we demonstrate how to build a rodent grip strength apparatus from scratch using a digital kitchen scale, an empty cage, and a microcontroller, with both hardware and software being completely open-source to enable maximal modularity and flexibility of the instrument in concordance with the principles of open-source bioinstrumentation. Furthermore, we test the griPASTA system for assessment of increased muscular rigidity in the proof-of-concept experiment in the rat model of Parkinson’s disease induced by intrastriatal administration of 6-hydroxydopamine (6-OHDA). Finally, the importance of bioinstrumental customization is demonstrated by utilizing griPASTA for assessment of trial speed from initial-to-maximal deflection time segments and controlling for its potential confounding effects on the grip strength.

**Significance Statement:** Neuromuscular function analyzed by grip strength performance tests is an integral part of motor system neuroscience and neurotoxicology. Strain gauge-based grip strength meters provide the most reliable results, however, commercial solutions are unaffordable by many. Consequently, cheap semi-quantitative tests are often used at the expense of precision and reliability. We propose griPASTA – a simple and robust open-source grip strength platform made from an ordinary kitchen scale. griPASTA could improve the quality and reproducibility of grip strength experiments by enabling researchers to obtain quantitative grip strength data in high resolution using a highly customizable platform.

## Introduction

Assessment of muscle strength is critical for understanding neuromuscular function in rodents. As many rodent disease models are characterized by impaired neuromuscular function, sensitive methods for objective quantification of muscle strength are a topic of interest. Numerous tests have been proposed for *in vivo* muscle strength assessment such as inverted screen (Kondziella, 1964), scale collector weight lifting (Deacon, 2013), wire hang (Hoffman and Winder, 2016), treadmill (Castro and Kuang, 2017), vertical pole (Matsuura et al., 1997), swimming endurance and grip strength test (Takeshita et al., 2017). The grip strength test is favored by most as it is quick, convenient, and introduces less stress than other tests (Takeshita et al., 2017). Its importance has been recognized by authorities - the test is included both in the US Environmental Protection Agency Functional Observational Battery (US EPA FOB) and OECD guidelines for neurobehavioral toxicity screening (Moser, 2011). Several commercial strain gauge-based grip strength meters for precise and objective quantification are available on the market, however, they are unaffordable for many. Consequently, researchers usually opt out for cheap semi-quantitative solutions for grip performance evaluation. We designed a simple open-source platform that combines affordability, availability, and simplicity of semi-quantitative grip strength tests, and precision, robustness and informativeness of the quantitative strain gauge-based methods.

Commercially available equipment for measurement of grip strength in rodents exploits strain gauges, sensors that convert force into a change in electrical resistance that can be easily measured. The same force-to-resistance conversion method is unknowingly used by many every day as cheap commercial kitchen scales exploit the very same principle for weight estimation. We previously proposed a method for the assessment of acoustic startle response and prepulse inhibition sensorimotor gating phenomenon in rodents that exploits the same principles (Virag et al., 2021). Based on the same open-source platform we present griPASTA – a solution for quantification of grip strength in experimental animals with a step-by-step guide on how to build it by transforming a kitchen scale.

We deem that cheap open-source solutions such as griPASTA might be particularly valuable in specific sensitive experimental settings where the modularity of the open-source devices enables the researchers to address specific methodological problems that would otherwise remain overlooked due to the limited flexibility of the black box solutions. For this reason, we evaluated ability of griPASTA to provide raw data to test for the potential confounding effect of the trial speed in our real-world proof-of-concept (POC) experiment in a rat model of Parkinson’s disease (PD) induced by intrastriatal administration of 6-hydroxydopamine (6-OHDA). This variable would remain largely ignored if a commercial apparatus was used for the grip strength assessment, although previous research identified trial speed as a factor that might affect the grip strength measurements (Maurissen et al., 2003). Muscular rigidity has been considered an important parameter in animal models of PD, however, it remains underrepresented in the literature due to the challenging methodology associated with rigidity assessment in rodents (Meredith and Kang, 2006). Different tests have been proposed for the purpose, such as the lateral displacement test (Lindner et al., 1999). Still, researchers often opt for other tests more closely related to the grip strength, such as the grasping test, for their better discriminatory power (Ma et al., 2014; Sotnikova et al., 2005). We hypothesized that quantitative grip strength assessment with griPASTA might be able to detect an increment in muscular rigidity in 6-OHDA-treated rats although previous reports indicated that intrastriatal lesions induce much less pronounced rigidity in comparison with nigrostriatal ones (Jeyasingham et al., 2001). Additional advantages of quantitative grip strength assessment by griPASTA over the grasping test include control for the confounding effect of body weight, body weight-grasping outcome linearity, cut-off determination, and ceiling effects (Liu and Wang, 2021).

## Materials and Methods

### Construction of griPASTA Hardware

A setup for the measurement of grip strength in rodents consists of two parts: a hacked digital kitchen scale connected to a computer (Virag et al., 2021) and a weighted mouse cage used as a grip test platform (**Fig 1A-C**).

**Fig 1.**
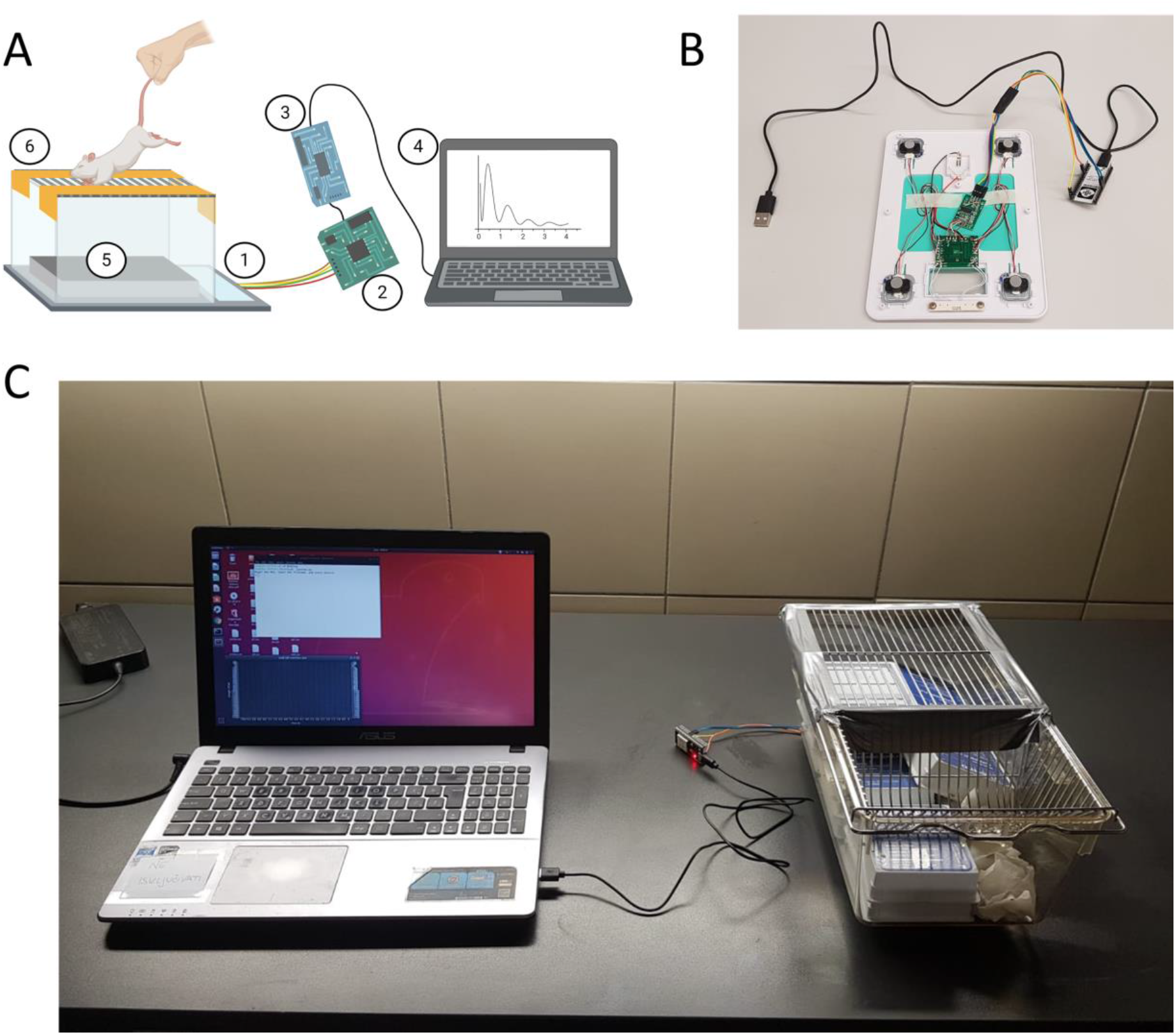
Concept of griPASTA. **A)** Schematic representation of griPASTA setup for grip strength performance measurements in rodents. A1 - PASTA: modified kitchen scale described by Virag *et al.* (Virag et al., 2021); A2 - HX711 circuit board; A3 - NodeMCU ESP-32S microcontroller; A4 - laptop; A5 - a weight stabilizing the mouse cage; A6 - Duct tape modified cage lid used as a grip platform. **B)** PASTA setup. A NodeMCU ESP-32S microcontroller interfaced with kitchen scale strain gauge load cells through the HX711 circuit board. **C)** A Photograph of griPASTA setup with a grip test platform fixed on top of the PASTA platform.

### A hacked kitchen scale

For detailed instructions on how to hack a kitchen scale and connect it to a computer as well as for free open-source software packages for data extraction and visualization see Virag et al.(Virag et al., 2021). In brief, a kitchen scale is opened and strain gauge load cells are identified and connected to a readily available HX711 circuit board modified by reconnecting the RATE pin to VCC to enable 80 samples/second sampling rate mode of the integrated circuit. The HX711 is connected to a microcontroller that acts as a communication bridge between the kitchen scale and the computer through a USB connection. We used a NodeMCU ESP-32S board, but other Arduino microcontroller boards may be used just as well, including AVR-based ones.

### A grip test platform

To extract grip strength data from a kitchen scale, we constructed a grip platform for rats. We wanted to use something readily available in most laboratories equipped for experiments with laboratory animals, so we used a mouse cage with a cage lid used as a test surface and several boxes of old histological slides as a weight to stabilize the platform. To reduce the variability of animal gripping patterns, duct tape was placed on the cage lid making only central grids available for the animals to hold onto. The whole system was fixed onto the PASTA platform with reusable putty-like pressure-sensitive adhesive (eg. Patafix, UHU, Germany).

### griPASTA Software

To extract data from the platform, we used a custom Python script written for general purpose experimentation with PASTA called PASTAgp (PASTA general-purpose script), with a modified Y scale for grip testing. Once loaded, the script asks for the ID of the animal and initiates a Python PyQtGraph plugin-based real-time plot of the scale-extracted data (**Fig 1C**). When the recording is done, the script automatically saves data in a comma-separated value format in the predefined directory, using the “.pasta” extension (ID.pasta) to simplify the loading of files in the downstream analysis. The whole code is freely available under the GPLv3 copyleft license in the PASTAgp GitHub repository (“GitHub - davorvr/pasta-gp,” n.d.).

### Development of a semi-automatic analysis algorithm and implementation in the ratPASTA R package

A semi-automatic function for easier analysis of data exported from griPASTA was developed and implemented in the ratPASTA package (Kodvanj et al., 2020) (available from CRAN and GitHub). Briefly, four functions were introduced: *loadgriPASTA*, *errors*, *findPeaks*, and *griPASTA*. The *loadgriPASTA* function imports and handles .pasta data. The function *errors* is an artifact identifier and processor that calls the interactive plotly-type plot with time values depicted on the x, and griPASTA value on the y axis. Once the user specifies the coordinates of an artifact, the function removes an artifact, returns the processed plot, and asks whether other artifacts are present. The loop continues until there are no artifacts left. The function *findPeaks* takes the errors-processed data and asks the user for the number of peaks and the estimated length of each trial in seconds. Once the information is provided, the function scans the griPASTA data and returns values of peaks corresponding to the modulus of maximal negative deflection times (a proxy for grip strength). Both *errors* and *findPeaks* are subfunctions used by the main *griPASTA* function. *griPASTA* is essentially a wizard function that takes the griPASTA data, and exploits *errors* and *findPeaks* to guide the user through the analysis process and helps with automation of the manual data extraction from the plot.

### Animals

Three-month-old male Wistar rats from the Department of Pharmacology, University of Zagreb School of Medicine were used in the experiment. Thirty-six rats weighing 368 (SD = 34 g) were randomly assigned to four groups (Control [CTR], 6-OHDA 6 μg [OHDA 6], 6-OHDA 6 μg w/o reboxetine, a noradrenaline transporter inhibitor [OHDA 6 w/o REB], and 6-OHDA 12 μg [OHDA 12]; n = 9 per group). The rats were housed 2-3 per cage under a 12-hour light cycle (7AM/7PM), at 21-23°C, with humidity in the 40-70% range. Standardized food pellets and water were available *ad libitum*. Standard wood-chip bedding was changed twice a week.

### Drug treatment protocol

6-OHDA was dissolved in 0.02% ascorbic acid in normal saline, and Reboxetine (Edronax, Pfizer) was dissolved in normal saline. All animals except the ones from the 6 OHDA w/o REB group received an intraperitoneal injection of reboxetine (5 mg/kg in 1ml) 40 minutes before induction of general anesthesia (intraperitoneal ketamine (70 mg/kg) / xylazine (7 mg/kg)). The skin was surgically opened and the skull was trepanated bilaterally with a high-speed drill. Intrastriatal administration of 6-OHDA solution (2 μl corresponding to 6 μg 6-OHDA /hemisphere; 2 μl corresponding to 12 μg 6-OHDA /hemisphere) or vehicle was done at stereotaxic coordinates of 0 mm posterior, ±2.8 mm lateral and 7 mm depth from pia mater relative to bregma. One animal in each group died during the induction procedure or within the following 24 hours, and two additional animals died in the group treated with the high dose of 6-OHDA within the following 2 weeks. Grip strength assessment with griPASTA was conducted 1 month after intrastriatal administration.

### Experimental protocol for grip strength assessment with griPASTA platform

A modified kitchen scale was placed on a clean and flat surface and connected to a computer via USB. A grip test platform was prepared by placing a weight in a clean mouse cage. In this study, a 2184L Eurostandard Type II L cage with dimensions of 365×207×140 mm was used. A cage lid, used as a grip platform was a grid of parallel wires with a diameter of 2 mm, and a 7 mm wide gap, and a single perpendicular wire with a diameter of 2.5 mm. The precise quantification range of the kitchen scale used (Vivax Home KS-502T (Vivax, Croatia)) was 0-5000 g so a platform weighing 4500g was created to maximally reduce the possibility to move during the testing procedure and to make sure that the negative force resulting from rat holding onto the cage lid while being removed by the experimenter would remain in the range of maximal quantification. When the platform was ready the PASTAgp script was loaded. Once the animal ID was provided, a PyQtGraph real-time plot of the scale-extracted data was initiated and the platform was calibrated by pressing the reset button on the microcontroller board (“EN” for NodeMCU ESP-32S). The animal was placed on the exposed part of the cage lid and elevated vertically by the tail. The negative force produced by animal resisting removal from the platform was recorded. Once the trial was finished, the real-time plot was closed and data was automatically saved in the .pasta file in the working directory.

In the 6-OHDA POC experiment, 5±1 consecutive measurements were recorded per trial for each animal.

### Animal body weight assessment

On the day of experiment, each animal was placed on a digital scale, and the bodyweight was read from the screen once the animal stopped moving.

### Data analysis and visualization

Data analysis and visualization were conducted in R 4.0.5. griPASTA (.pasta) data were imported into R and moduli of maximal negative deflection values and temporal values of maximal and initial deflection points for all trials were extracted manually from interactive plots created by Plotly open-source JavaScript graphing library. Grip strength differences between groups, the potential effect of animal weight and trial speed were examined by linear mixed-effects models (LMM) with the restricted maximum likelihood estimation and animal ID used as a random effect. Holm correction was used to control the familywise type 1 error rate. Alpha was set at 5%.

### Ethics committee approval

Animal procedures were carried out at the University of Zagreb Medical School (Zagreb, Croatia) and complied with current institutional, national (The Animal Protection Act, NN102/17; NN 47/2011), and international (Directive 2010/63/EU) guidelines governing the use of experimental animals. The experiments were approved by the Ethical Committee of the University of Zagreb School of Medicine and the Croatian Ministry of Agriculture (229/2019).

## Results

### Exploration of griPASTA data and semi-automatic extraction using ratPASTA R package

Exploratory analysis of griPASTA data from a single trial confirmed the hypothesis that load cells from the modified PASTA platform (Virag et al., 2021) can be used to analyze different aspects of rodent motor behavior. Raw data from the trial (**Fig 2A**) and a single measurement (**Fig 2B**) reveal several informative patterns. Placement of the animal on top of the platform generated a positive deflection from baseline with maximal values approximately corresponding to animal weight (**Fig 2B-2**). Although analysis of the positive deflection could theoretically be used for bodyweight assessment, in this particular case the data were not sufficiently reliable due to the animal moving on top of the platform, and the experimenter holding the animal by the tail. Once the experimenter tries to remove the animal from the platform a negative deflection is recorded. Maximal negative deflection values correspond to the peak force the animal was able to resist, and the modulus of this value was considered an indicator of grip strength (**Fig 2B-3**). The time between the first deflection from the plateau and the moment when maximal deflection value has been achieved (initial-to-maximal deflection time segment) was considered a proxy of trial speed – i.e. how fast the experimenter attempted to remove the animal from the platform (a blue segment in **Fig 2B**).

**Fig 2.**
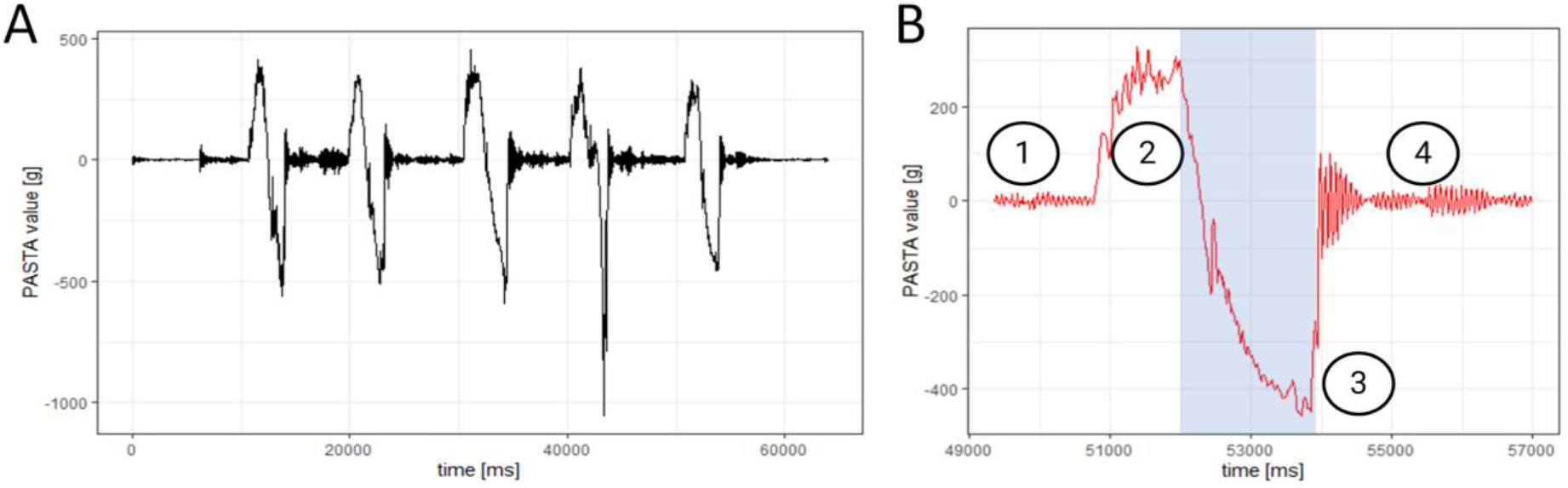
**A)** Raw data from a single trial with 5 consecutive measurements. **B)** An example of raw data from a single measurement. B1 - baseline data; B2 - positive deflection caused by the placement of the animal on top of the platform (positive deflection indicates the weight of the animal); B3 - negative deflection caused by animal holding onto the platform while being removed by the tail. The maximal negative value is the peak force the animal was able to resist; A modulus |X| of this value (|B3|) was taken as an indicator of grip strength; B4 - baseline values with noise recorded due to the placement of the animal on the table next to griPASTA. The initial-to-maximal deflection time segment, used as a proxy for trial speed, is indicated in blue.

The process of griPASTA data extraction was simplified by introducing 4 griPASTA functions in the ratPASTA package: *loadgriPASTA, errors, findPeaks,* and *griPASTA* function with the latter being the most important function for non-experienced users (Kodvanj et al., 2020). An example of griPASTA-guided extraction of grip strength from a single animal trial is shown in **Fig 3**.

**Fig 3.**
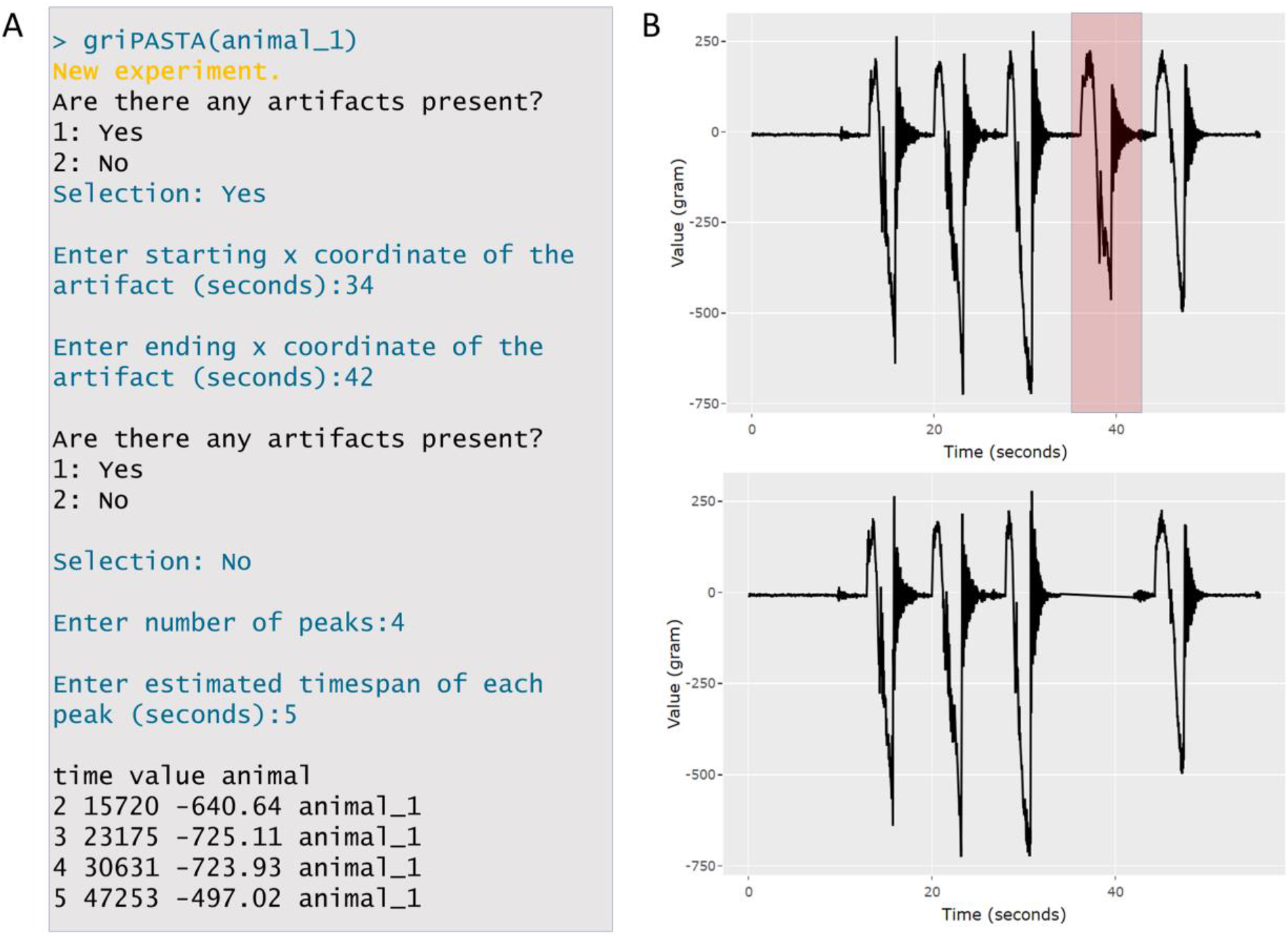
An example of the *griPASTA*-assisted grip strength data extraction from the raw griPASTA-obtained data in R. **A)** An example of the user-griPASTA wizard interaction from the R console. **B)** A plotly-type interactive plot called by the *griPASTA* function. For demonstrative purposes, the fourth experimental measurement (highlighted in red) from the trial was used as an artifact since the animal didn’t hold onto the grip properly. The trial was marked as an artifact and removed by the *griPASTA* wizard using the *errors* function. The rest of the peaks were automatically extracted by the *findPeaks* and reported to the user in the console (A).

### Grip strength in rats following intrastriatal administration of two different doses of 6-OHDA

Grip strength of 6-OHDA-treated animals and respective controls was assessed using the same protocol. Raw data corresponding to moduli of maximal negative deflection values, speed of each trial inferred from initial-to-maximal deflection time segments, and weight of each animal were reported (**Fig 4A**). Mean averages of all measurements from a single trial were reported separately (**Fig 4B**). As evident from the individual (**Fig 4A**) and pooled measurements (**Fig 4B**), and the models (**Table 1**), 6-OHDA-treated rats display forelimb motor rigidity. This is in concordance with previous findings in 6-OHDA treated rats that demonstrate increased hang time in the hanging grip strength test as compared to controls (Ma et al., 2014). Grip strength assessment is known to be under influence of numerous factors that are often overlooked (e.g. animal weight and trial speed (Maurissen et al., 2003; Takeshita et al., 2017)). As the ability to access raw data enables measuring such potential confounders, we examined whether the differences of grip strength estimates obtained by griPASTA would still be present after accounting for trial speed and animal weight. In this particular experiment, neither animal weight nor trial speed significantly affected grip strength measurements (**Table 1.**).

**Table 1.** griPASTA models. Linear mixed models of griPASTA taking into account only treatment (Model 1), treatment and animal weight (Model 2), treatment and trial speed (Model 3), and treatment, animal weight, and trial speed (Model 4). Animal ID was included as a random effect in all models.

|  | Model 1 |  |  | Model 2 |  |  | Model 3 |  |  | Model 4 |  |  |
| --- | --- | --- | --- | --- | --- | --- | --- | --- | --- | --- | --- | --- |
| Predictors | Estimates | CI | p | Estimates | CI | p | Estimates | CI | p | Estimates | CI | p |
| (Intercept) | 542.16 | 430.25 – 654.06 | <.001 | 542.63 | 425.82 – 659.43 | <.001 | 536.15 | 422.43 – 649.86 | <.001 | 536.63 | 418.06 – 655.20 | <.001 |
| OHDA 12 vs CTR | 230.50 | 60.43 – 400.58 | 0.008 | 229.09 | 43.01 – 415.16 | 0.032 | 242.87 | 67.89 – 417.86 | 0.013 | 241.47 | 50.80 – 432.13 | 0.039 |
| OHDA 6 vs CTR | 261.10 | 102.58 – 419.62 | 0.004 | 260.29 | 92.30 – 428.28 | 0.007 | 266.06 | 106.60 – 425.53 | 0.004 | 265.26 | 96.35 – 434.18 | 0.008 |
| OHDA 6 w/o REB vs CTR | 260.77 | 102.51 – 419.03 | 0.004 | 260.93 | 99.36 – 422.51 | 0.006 | 269.82 | 108.66 – 430.98 | 0.004 | 269.96 | 105.51 – 434.40 | 0.006 |
| weight | - | - | - | -0.02 | -1.41 – 1.37 | 0.976 | - | - | - | -0.02 | -1.41 – 1.37 | 0.977 |
| trial speed | - | - | - | - | - | - | -0.02 | -0.09 – 0.05 | 0.553 | -0.02 | -0.09 – 0.05 | 0.555 |

**Fig 4.**
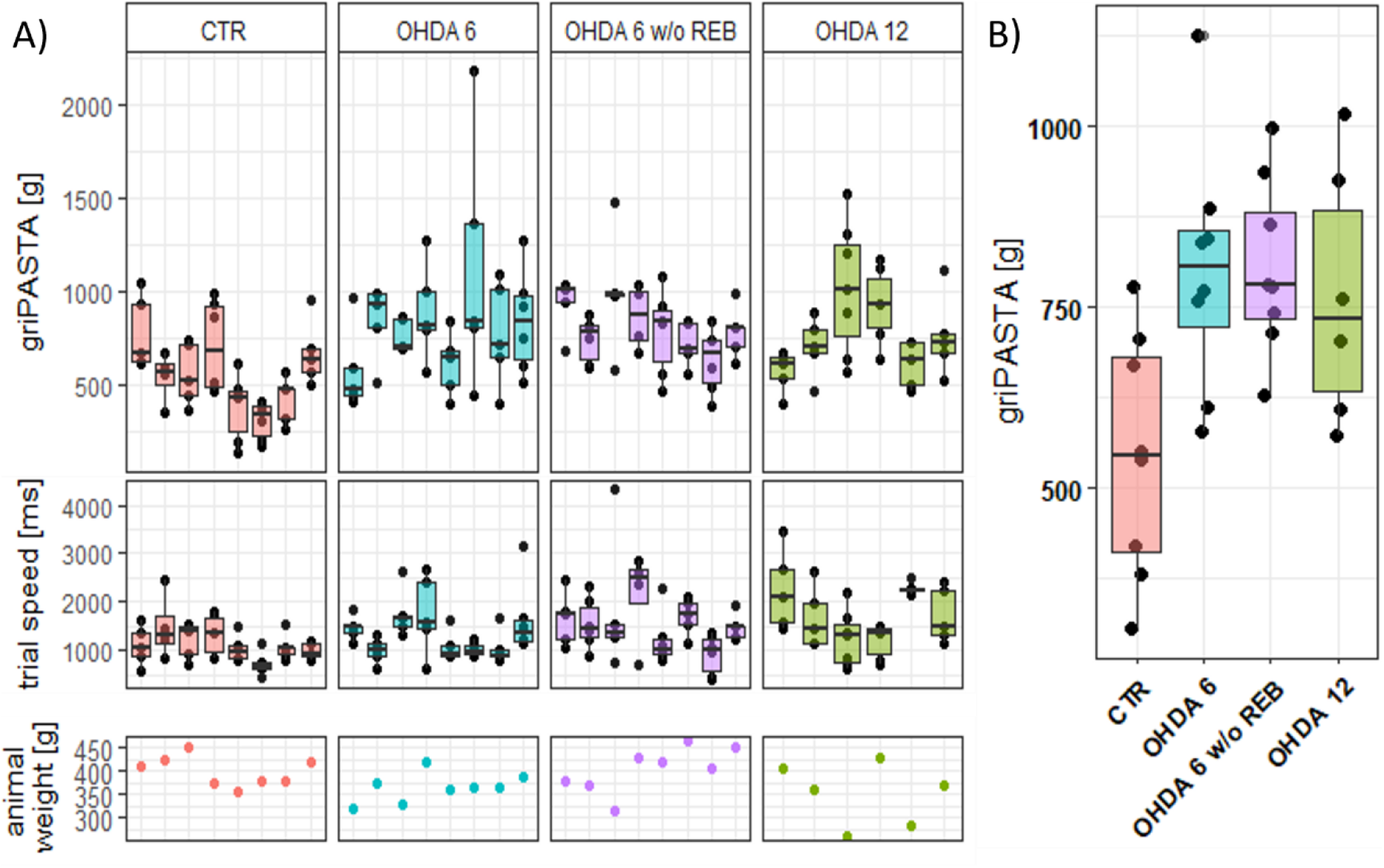
The effect of intrastriatal 6-OHDA treatment on the forelimb rigidity of rats analyzed by griPASTA grip strength assessment. **A)** Boxplot of moduli of maximal negative deflection values with data points corresponding to individual intra-trial measurements for each animal, speed of each trial inferred from initial-to-maximal deflection time segments, and the weight of each animal. **B)** Group differences displayed by a group boxplot with data points corresponding to mean averages of trial measurements for each animal. CTR – control animals; OHDA 6 – animals treated with 6 μg 6-hydroxydopamine/hemisphere; OHDA 6 w/o REB – animals treated with 6 μg 6-hydroxydopamine/hemisphere without reboxetine pretreatment; OHDA 12 – animals treated with 12 μg 6-hydroxydopamine/hemisphere.

## Discussion

Custom-made instruments have been used over the years to obtain continuous level grip strength data. In 1979, Meyer *et al.* proposed an affordable, rapid, and efficient method for forelimb and hindlimb grip strength assessment in rats and mice based on a mechanical strain gauge (Meyer et al., 1979). The proposed method was modified and used for toxicological research by different groups. For example, Mattsson *et al.* used a custom-built device based on a horizontal coarse wire screen attached to a mechanical strain gauge with a clutch which retains the position of the maximal reading in their analysis of neuropathic consequences of repeated dermal exposure of 2,4-dichlorophenoxyacetic acid (Mattsson et al., 1986). Development and increased availability of electronics led to increased use of electronic components in custom-made instruments based on the original proposal by Meyer *et al.* (Meyer et al., 1979). In an important study on the factors affecting grip strength testing by Maurissen et al., a custom-built electronic version of the original grip strength apparatus was used (Maurissen et al., 2003). The authors built a device based on the Chatillon DFA series digital force gauge (DFA-R-ND with F31197 10 kg load cell (AMETEK, USA)) connected to a 7 mm^2^ wire grid with corner support and compared the performance of the custom-built apparatus to the commercial grip strength measurement system 1027DSx (Columbus Instruments, USA). Although custom made instruments for grip strength assessment in rodents have been described in the literature, open-source solutions that would enable researchers to easily implement simple devices into their laboratory practice, use the flexibility of the freely available hardware and software to best adapt the system for their specific experimental goals, are scarce. One example is a bilateral grip strength assessment device (Kivitz and Lake, 2017) used for grip analysis in a rat model of post-traumatic elbow contracture (Reiter et al., 2019). The importance of modularity of the grip strength performance system is further corroborated in the study by Maurissen *et al.* (Maurissen et al., 2003). The researchers bought a commercial instrument, but also built their own, with the main motivation being the concern that sampling rate and platform support, both fixed and unmodifiable in the commercial setup, might significantly affect the outcome of the grip strength measurements (Maurissen et al., 2003).

We describe a simple, low-cost, and robust system for quantitative assessment of grip strength performance in rodents based on an open-source hacked digital kitchen scale multimodal platform called PASTA (Virag et al., 2021). The idea behind griPASTA is to record the force exerted by an animal resisting experimenter who is trying to remove it by the tail from the grip test platform placed on top of the PASTA system. The platform records the force with high temporal resolution and stores the unprocessed data, enabling flexibility in the process of subsequent data analysis. Consequently, the griPASTA platform can provide more information than most of the expensive commercial products used for this purpose. Furthermore, in contrast to commercial equipment that is usually relatively static in terms of design, griPASTA is characterized by simple and modular hardware based on readily available equipment in the animal research setting. In further text, we briefly discuss advantages of griPASTA open-source hardware and software in the context of grip strength assessment in rodents, with a special focus on trial speed and grip strength testing angle.

### Trial speed

Commercial instruments standardly measure maximal force the animal can resist while being removed by the tail from the test platform, but do not provide information on trial speed – a potential confounding factor that might influence grip strength assessment in rodents (Maurissen et al., 2003; Takeshita et al., 2017). Consequently, differences in trial speed between groups that arise either due to biological differences or by chance due to poor research design cannot be perceived nor accounted for. In contrast, assessment of initial-to-maximal deflection time segments from raw griPASTA data provides an insight into trial speed, enables its quantification, and subsequent control for its effect by statistical modeling. This is particularly important as there were few attempts to assess the influence of trial speed on the grip strength performance (e.g. (Maurissen et al., 2003)), and the results from „healthy” animals (Maurissen et al., 2003) do not take into account the potential differences that could arise due to pathophysiological processes in animal models of diseases. Consequently, griPASTA could improve the overall quality and reproducibility of grip strength experiments by empowering researchers to more easily detect and control for bias.

### Trial angle

Standard rodent grip strength assessment protocols are based on horizontal systems where the animal is pulled backward by the experimenter after it grasps the grip strength meter platform (Aartsma-Rus and van Putten, 2014; Cabe et al., 1978). The angle at which the animal is pulled away from the platform affects the overall grip strength assessment (Maurissen et al., 2003; Takeshita et al., 2017). Takeshita *et al.* proposed a simple modification based on the vertical rotation of the platform to improve the reliability and consistency of the conventional method (Takeshita et al., 2017). Vertical rotation of the platform ensures the animals are maximally motivated to keep grasping the bar while being pulled resulting in lower inter-trial and inter-day variability in comparison with the conventional protocol. The optimal position of the platform depends on the sex and age of the animals (Takeshita et al., 2017) and possibly other factors in animal models of diseases. For this reason, platform angle is usually flexible in the commercial setups and can be set at a desirable angle in the predefined range (eg. (“SDI motor & sensory test systems,” 2018)). The one-size-fits-all principle cannot be applied to the grip strength assessment. Consequently, maximal modularity of the design of a grip strength apparatus is important as it enables the researchers to adapt the instrument for a specific experimental purpose. griPASTA platform ensures maximal flexibility. In the present POC experiment, we used a horizontal grip platform made from a rodent cage lid (**Fig 1**). Nevertheless, the platform can be modified in various ways. A horizontal pull preferred by some laboratories can be measured by either constructing a holder for the apparatus or introducing a simple pulley (**Fig 5A**). Furthermore, a vertical setup, as proposed by Takeshita *et al.* (Takeshita et al., 2017), can be created simply by placing the whole griPASTA platform on the holder (**Fig 5B**). Additionally, griPASTA can be connected to a cheap and widely available load cell fixed permanently in the vertical setup (**Fig 5C**), and the instrument can be plugged in either a kitchen scale-based platform or in a vertically positioned strain gauge depending on the requirements of a specific experiment.

**Fig 5.**
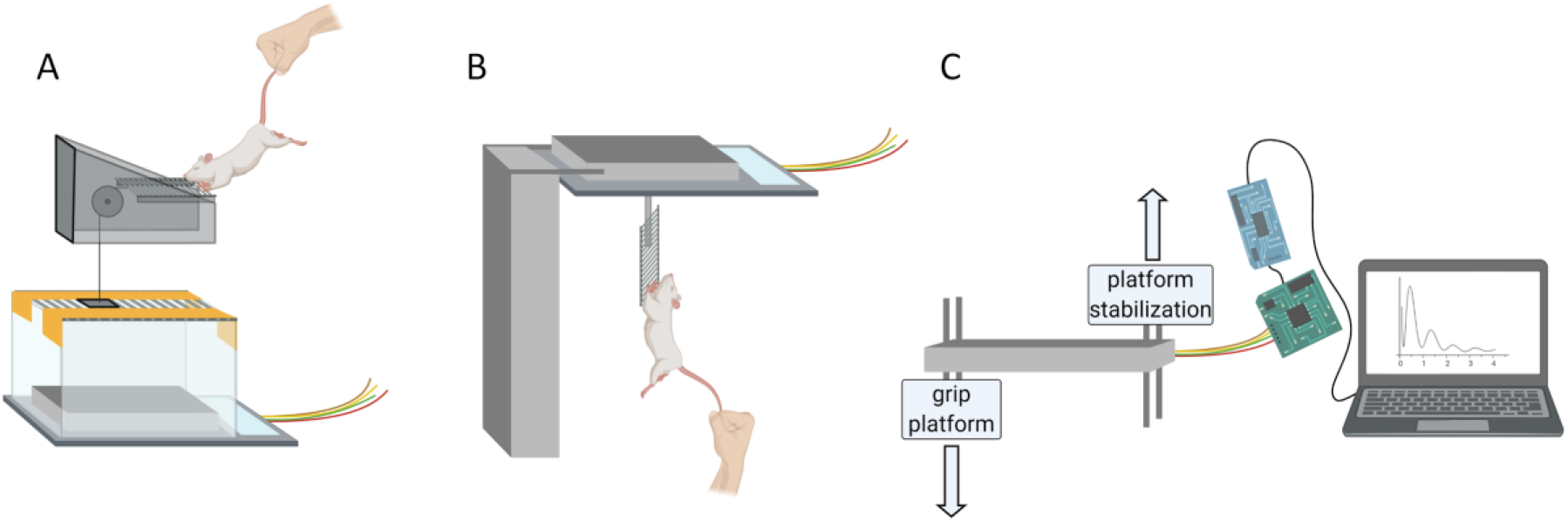
A schematic representation of the modular design of griPASTA adapted for different grip strength measurement strategies. **A)** A variant of the griPASTA grip strength assessment with pulley-based force transduction to ensure the horizontal pull. **B)** A vertical griPASTA setup, as proposed by Takeshita *et al.*, with a custom holder used as a griPASTA platform to align the load cell displacement vector with the pulling motion and ensure maximal motivation of the animal to resist being removed from the platform. **C)** For the easier implementation of the vertical setup, the introduction of an additional load cell is proposed, so the experimenter can simply plug in the griPASTA system either in a standard kitchen scale-based setup or in a single load cell-based vertical setup with minimal modifications.

### Grip strength as an indicator of muscular rigidity in rat models of PD

Alongside akinesia, bradykinesia, dystonia, resting tremor, and gait abnormalities, muscular rigidity is an important symptom of PD, and an important feature of successful animal models of PD (Meredith and Kang, 2006; Potashkin et al., 2010). Muscular rigidity, characterized by increased resistance to passive movement, has been proposed to emerge as a result of potentiation of long-latency electromyographic components related to the stretch reflex and co-activation of antagonistic muscles (Wolfarth et al., 1996). Among other animal models of PD, it has been suggested that 6-OHDA treated rats successfully mimic Parkinson-like muscular rigidity (Cenci et al., 2002; Wolfarth et al., 1996). Wolfarth *et al.* reported that bilateral administration of 1ul of 6.5 μg/μl 6-OHDA into substantia nigra induces muscle rigidity reflected both by mechanomyography and movement-induced reflex electromyographic activity (Wolfarth et al., 1996), and similar results have been reported following intrastriatal administration of 4 μg of 6-OHDA (Klockgether et al., 1987). Quantification of rodent muscular rigidity has been considered a daunting and challenging task and many different assays have been proposed over the years (Meredith and Kang, 2006)(e.g. the forced lateral movement test (Lindner et al., 1999)). A grasping test has also been used for this purpose (Jolicoeur et al., 1991). For example, Sotnikova *et al.* reported increased rigidity in dopamine transporter knockout mouse model of PD measured by the grasping test (Sotnikova et al., 2005), and Ma *et al.* reported increased rigidity by the same method in the medial forebrain bundle 6-OHDA-treated rats (Ma et al., 2014). Being closely related to the grasping test, some authors proposed that the grip strength test might also reflect rigidity in the animal models of PD. A marked increase in grip strength has been observed upon administration of 6-OHDA into the nigrostriatal bundle (Dunnett et al., 1998; Jeyasingham et al., 2001). Although the mechanism remains to be elucidated, it has been proposed that it reflects processes related to nigrostriatal degeneration-induced rigidity in human PD (Dunnett et al., 1998). Here, we exploited the griPASTA system to assess muscular rigidity following bilateral intrastriatal 6-OHDA administration. Intrastriatal lesions induce much less pronounced muscular rigidity in comparison with nigrostriatal lesions (Jeyasingham et al., 2001), however, we hypothesized that the quantitative nature and sensitivity of griPASTA might be able to detect the differences in motor rigidity we noticed while handling animals. The results of the POC experiment indicate that 6-OHDA, regardless of the dose or reboxetine pretreatment, significantly increases motor rigidity in rats (**Fig 4**, **Table 1**), and that muscular rigidity in rodent models of PD usually examined by grasping tests might also be assessed using quantitative grip strength testing. Taking into account the limitations of the standard grasping tests, such as the effect of body weight, and the problem of its linearity, as well as the choice of the cut-off time and subsequent statistical problems related to the ceiling effects (Liu and Wang, 2021), the grip strength test should be explored as an alternative solution for quantification of muscular rigidity in this context. The latter is especially true considering that open-source low-cost solutions such as griPASTA enable quantification without sophisticated and expensive research equipment. Finally, our results confirm previous findings of muscular rigidity as a pathological hallmark of 6-OHDA-lesioned rats and encourage further investigation of this marginalized symptom in animal PD research.

The griPASTA project is fully open-source and is described in concordance with the proposed principles of open-source bioinstrumentation (Booeshaghi et al., 2019), gaining considerable attention in recent years (Maia Chagas, 2018; Pearce, 2014, 2012; White et al., 2019). As already emphasized, apart from significant economic advantages of open-source lab equipment (Dolgin, 2018), the main strength of open-source bioinstrumentation stems from modularity, flexibility, and the active engagement of the scientific community to implement and further improve freely available hardware and software solutions (Booeshaghi et al., 2019). We consider griPASTA to be a work in progress, a project we remain actively engaged with, and use this opportunity to invite our colleague scientists to offer their opinions and suggestions and become actively involved in its further development.

## Conclusion

griPASTA is a simple open-source PASTA-based (Virag et al., 2021) solution for grip strength assessment in rodents that can be made on the cheap using readily available utensils. The whole setup consists of a hacked kitchen scale connected to a computer and a mouse cage used as a grip platform. Apart from the cost, griPASTA provides several advantages over the commercially available solutions as the platform provides raw data which makes it compatible with different data analysis strategies and allows the researcher to control for potential confounding factors such as trial speed. Additionally, griPASTA open-source hardware and design are maximally flexible which makes the platform highly customizable, so with minimal modifications both platform and load cell angle can be easily adapted. Finally, our POC experiment suggests that griPASTA provides a good insight into motor dysfunction and rigidity in the 6-OHDA rat model of Parkinson’s disease. Considering the observed differences, robustness, low cost, and simple design of the griPASTA platform, we believe that the proposed quantitative grip strength assessment protocol should be further explored in different animal models as it might provide important additional information on motor dysfunction rapidly and economically.

## Data and code availability statement

A general-purpose python script for extraction of data from the PASTA platform (PASTAgp) is available on GitHub at https://github.com/davorvr/pasta-gp

Data from the POC experimental trial (ID.pasta) and a video example of a single measurement trial are available on GitHub at https://github.com/janhomolak/gripasta

ratPASTA is available on CRAN and on GitHub (https://github.com/ikodvanj/ratPASTA).

## Author’s contributions

**JH** conceptualized the experiment. **JH, ABP, AK,** and **JOB** carried out drug treatments. **JH** constructed the griPASTA platform, **DV** contructed PASTA and wrote the PASTAgp script, and **IK** expanded the ratPASTA package with griPASTA functions. **JH** carried out the experiment, collected and curated the data. **JH** and **VT** analyzed data. **JH** visualized the data and wrote the manuscript. **DV**, **IK**, **IM**, **ABP**, **AK**, **JOB**, **VT,** and **MSP** commented on the manuscript and provided critical feedback. **MSP** and **IM** provided funding. **MSP** (mentor of JH and PI of the lab) supervised the project.

